# Glucose Metabolism echoes Long-Range Temporal Correlations in the Human Brain

**DOI:** 10.1101/2025.07.29.667370

**Authors:** Massimiliano Facca, Anna Ridolfo, Miriam Celli, Claudia Tarricone, Ilaria Mazzonetto, Tommaso Volpi, Andrei G. Vlassenko, Manu S. Goyal, Maurizio Corbetta, Alessandra Bertoldo

## Abstract

Intrinsic brain activity is characterized by pervasive long-range temporal correlations. While these scale-invariant dynamics are a fundamental hallmark of brain function, their implications for individual-level metabolic regulation remain poorly understood. Here, we address this gap by integrating resting-state functional Magnetic Resonance Imaging (fMRI) and dynamic [^18^F]FDG Positron Emission Tomography (PET) data acquired from the same cohort of participants. We uncover a systematic relationship between long-range temporal correlations, quantified via the Hurst exponent, and glucose metabolism. Our findings reveal that persistent temporal dependencies impose a measurable metabolic cost, with brains exhibiting higher long-range temporal correlations incurring greater energetic demands. Beyond glucose metabolism, we also show that these dynamics are likely supported by continuous biosynthetic processes, such as protein synthesis, which are critical for neural circuit maintenance and remodeling. Overall, our results suggest that a significant fraction of the brain’s so-called “Dark Energy” is actively spent to power spontaneous long-range temporal correlations.

## Introduction

The external environments and internal representations we humans navigate daily are structured across multiple, nested temporal scales. To make sense of the world – whether following a conversation or planning a future action – the brain must bridge milliseconds to minutes, weaving fragmented inputs into a coherent narrative (Baldassano et al. 2017; Hasson et al. 2015). This seamless integration is achieved through multiple processing streams, each operating with its own intrinsic temporal scale, or *Temporal Receptive Window* (TRW, Hasson et al. 2008; Lerner et al. 2011). These timescales, initially compressed in early sensory areas to match the rapid flow of input, progressively expand along the sensory-fugal axis of the brain’s organization (Murray et al. 2014; Raut et al. 2020). At the apex of this hierarchy, TRWs reach their maximal dilation, and activity slowly evolves while keeping track of its previous states – a phenomenon known as *long-range temporal correlations* (Hasson et al. 2008; He 2014; Linkenkaer-Hansen et al. 2001; Novikov et al. 1997; Raut et al. 2020). Temporal dependencies in neural activity are not only hierarchically organized, but also correlated between task and resting states (Golesorkhi et al. 2021; Ito et al. 2020), suggesting that their spatial arrangement is at least partially hardwired into the brain’s circuitry (Gao et al. 2020). Both computational and empirical studies indicate that highly-connected hubs in the human brain operate at slower timescales (Chaudhuri et al. 2015; Fallon et al. 2020; Gollo et al. 2015), enhancing integration and maintaining an efficient ebb and flow of information across the network - a principle that extends beyond the primate lineage (Sethi et al. 2017). Despite regional differences in intensity and patterning, sluggish temporal dependencies are a ubiquitous feature of the healthy human brain (He 2014; He et al. 2011). From a complex systems perspective, long-range temporal correlations are considered hallmarks of a system approaching criticality, i.e., the *critical slowing down (O’*Byrne & Jerbi 2022; Cocchi et al. 2017). Accordingly, in states that push the brain away from the critical point, such as loss of consciousness, these temporal dependencies have been shown to fade, only to rebound with its recovery (Barttfeld et al. 2015; Tagliazucchi et al. 2016; Tagliazucchi et al. 2013).

Like any other organ, the brain operates at an energetic cost. Its energy budget, however, greatly defies its size: accounting for just 2% of body weight, it consumes approximately 25% of the body’s energy (Clarke & Sokoloff, 1999). [^18^F]Fluorodeoxyglucose Positron Emission Tomography ([^18^F]FDG-PET) is currently the most effective tool to quantify glucose metabolism in the living human brain (Sokoloff et al. 1977). Decades of [^18^F]FDG-PET studies have shown that the metabolic cost of spontaneous brain activity far exceeds that of task-related processing (Attwell & Laughlin 2001; Raichle & Mintun 2006). This disproportionate energy use at rest has been metaphorically described as the brain’s “Dark Energy” (Raichle 2006). To map spontaneous neural activity, resting-state functional Magnetic Resonance Imaging (rs-fMRI) has long stood as the modality of choice, offering a favorable balance of spatial and temporal resolution (Fox & Raichle 2007; Van den Heuvel & Hulshoff Pol 2010). Multimodal studies integrating [^18^F]FDG-PET and rs-fMRI, however, have revealed only modest convergence: the network-based paradigm, dominant in rs-fMRI literature, accounts poorly for regional variability in glucose metabolism (Aiello et al. 2015; Riedl et al. 2014; Palombit et al. 2022). In contrast, local activity measures show a more consistent association with glucose consumption, suggesting that neuroenergetic coupling is more faithfully captured at the local rather than network level (Volpi et al. 2024; Volpi et al. 2025). In a similar vein, recent rs-fMRI research has shown that many network-level phenomena can be explained by local properties such as temporal autocorrelation, which may ultimately help link neural activity to its underlying biology (Shinn et al. 2023).

Motivated by these findings, here we systematically chart the metabolic consequences of long-range temporal correlations in the human brain. Our analysis is based on a multimodal dataset comprising rs-fMRI and dynamic [^18^F]FDG-PET data (Goyal et al. 2023). We hypothesize that the temporal persistence of brain activity, quantified by the Hurst exponent (Linkenkaer-Hansen et al. 2004; Maxim et al. 2005; Wink et al. 2008), is an energetically demanding process that contributes to glucose consumption at rest (Figure 1A). Adopting a multilevel perspective (Figure 1C), we show that: (i) regional variability in glucose consumption spatially mirrors regional increases and decreases in long-range temporal correlations at both group and individual levels, and (ii) brains exhibiting stronger long-range temporal correlations incur higher energetic costs than those with weaker correlations. We also exploit PET template maps, based on independent datasets, to show that the temporal stability of brain activity may be bestowed by a high-rate of protein synthesis and turnover (Schmidt et al. 2005). These findings suggest a link between intrinsic temporal dynamics and the brain’s metabolic/biosynthetic architecture. Collectively, our results indicate that a portion of the brain’s so-called “Dark Energy” fuels the long-term memory of neural dynamics.

**Figure 1.**
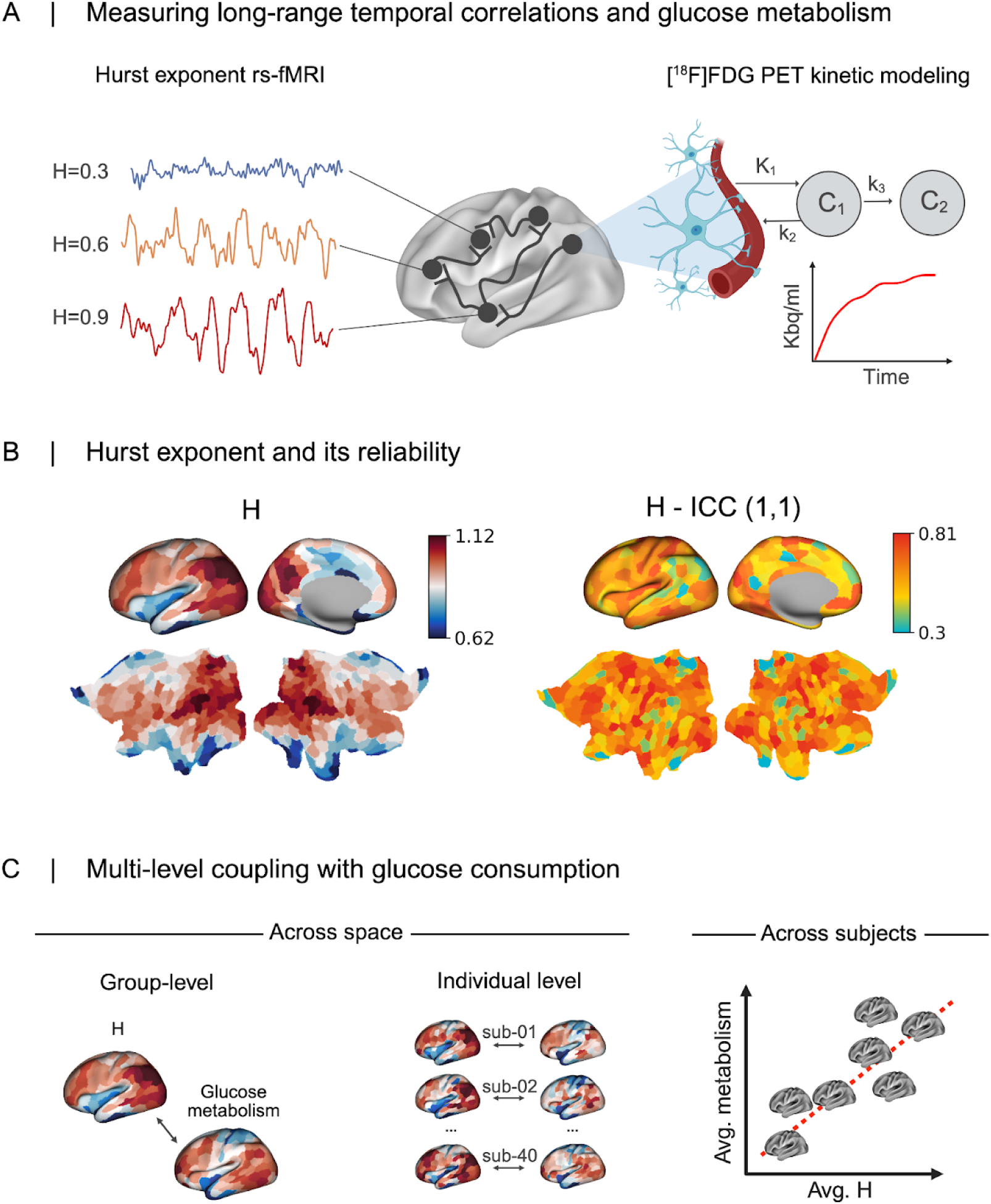
O**v**erview **of the study.** A) In this work long-range temporal correlations are measured as the Hurst exponent of regional rs-fMRI time series. Conversely, regional glucose metabolism is measured using dynamic [^18^F]FDG-PET in a quantitative fashion by means of kinetic modeling. B) Group template map of the Hurst exponent (H). Two patterns clearly emerge: 1) long-range temporal correlations are heterogeneously distributed across the brain and 2) the Hurst exponent is consistently greater than 0.5, indicating a ubiquitous persistence of brain activity. For subcortical structures see Supplementary Materials figure S1. In the right panel, the intraclass correlation coefficient (ICC-1,1) is shown to highlight the reliability of the Hurst exponent across different rs-fMRI sessions. C) Overview of the analytical approaches implemented in the present study. The coupling between glucose metabolism and long-range temporal correlations was investigated both across space, i.e., spatial correlations at the group and individual-subject levels, and across subjects, i.e., the correlation between the average Hurst exponent and average glucose metabolism across subjects.

## Results

This study is based on a cohort of 43 healthy subjects who underwent rs-fMRI and dynamic [^18^F]FDG-PET (Goyal et al. 2023). This allows us to investigate how coupled long-range temporal correlations and glucose metabolism are at both group and single-subject levels. Operationally, we measure long-range temporal correlations with the Hurst exponent (Hurst 1951). This metric has been widely employed for this purpose in both biological systems (Ciuciu et al. 2012; DePetrillo et al. 1999; Peng et al. 1994) and non-biological systems (e.g., Milanese et al. 2019; Shao & Ditlevsen 2016).

### Long-range temporal correlations in spontaneous brain activity

We begin with mapping the Hurst exponent distribution across the brain using a wavelet-based estimator (see Methods). We adopt a popular functional parcellation of the cerebral cortex in 400 regions (Schaefer et al. 2018), augmented with a subcortical atlas consisting of 16 regions (Tian et al. 2020). At first glance, we observe that long-range temporal correlations are ubiquitous in the brain: in every subject and every brain region, the Hurst exponent remains above H = 0.5. This value serves as a cut-off to identify persistent behavior. Although ubiquitous, long-range temporal correlations are unevenly distributed across regions. We observe the highest Hurst exponent values in the precuneus and the angular gyrus (Figure 1B). Since the study cohort consists of multiple sessions of rs-fMRI, we evaluate the extent to which the Hurst exponent is stable across sessions. We evaluate stability as the Intraclass Correlation Coefficient (ICC-1,1 McGraw & Wong, 1996). Conventionally, an ICC value greater than 0.4 indicates a good across-session reliability (Cicchetti 1994). In our data, this criterion is met in nearly all brain regions, with a median ICC of 0.61 (MAD = 0.07). The high test-retest reliability of the Hurst exponent across sessions suggests that long-range temporal correlations are a stable feature of brain activity.

### The costly architecture of spontaneous long-range temporal correlations

Glucose metabolism in this work is inferred from dynamic [^18^F]FDG-PET. Crucially, here we exploit a full kinetic model of [^18^F]FDG which dramatically enhances its biological insights (Schmidt & Turkheimer 2002). The model allows us to disentangle three key steps of the [^18^F]FDG kinetics in the brain: influx across the blood-brain barrier (BBB), clearance or efflux, and phosphorylation catalyzed by hexokinase, i.e., the actual utilization (Sokoloff et al. 1977). Each step is associated with a constant: *K_1_* (influx, [mL/cm^3^/min]), *k*_2_ (efflux, [min^-1^]) and *k_3_* (phosphorylation, [min^-1^]). These three are called micro-parameters describing tracer kinetics. Additionally, we consider the *irreversible uptake rate* (*K_i_*, [mL/cm^3^/min]) which is a macro-parameter that accounts for all the micro-parameters (Sokoloff et al. 1977). We generate a group template map for each parameter and assess the spatial association with the Hurst exponent (Figure 2A,B). A similar regional distribution between two measures constitutes a first indication that they are coupled. To ensure that our results are not driven by spatial autocorrelation, we employ an appropriate null model for significance testing (Burt et al. 2020). The group-level regional distribution of the Hurst exponent is strongly and positively correlated with *K_i_* (r = 0.65, p_SMASH_ < 0.0001). Examining the correlation with the micro-parameters, we observe that k_3_ exhibits the strongest relationship with the Hurst exponent (r = 0.44, p_SMASH_ < 0.0001), whereas *K_1_* and *k_2_*, which index [^18^F]FDG delivery and efflux, respectively, show no statistically significant association (Figure 2B and Supplementary Materials, Figure S3). This nourishes the idea of a genuine link between glucose’s *actual* utilization and long-range temporal correlations. Lastly, we ensure that the correlational patterns we have found are also replicable at the individual level: the correlational patterns broadly recapitulate group-level data (Figure 2C). Collectively, brain regions characterized by stronger long-range temporal correlations utilize more glucose than regions with weaker temporal dependencies.

**Figure 2.**
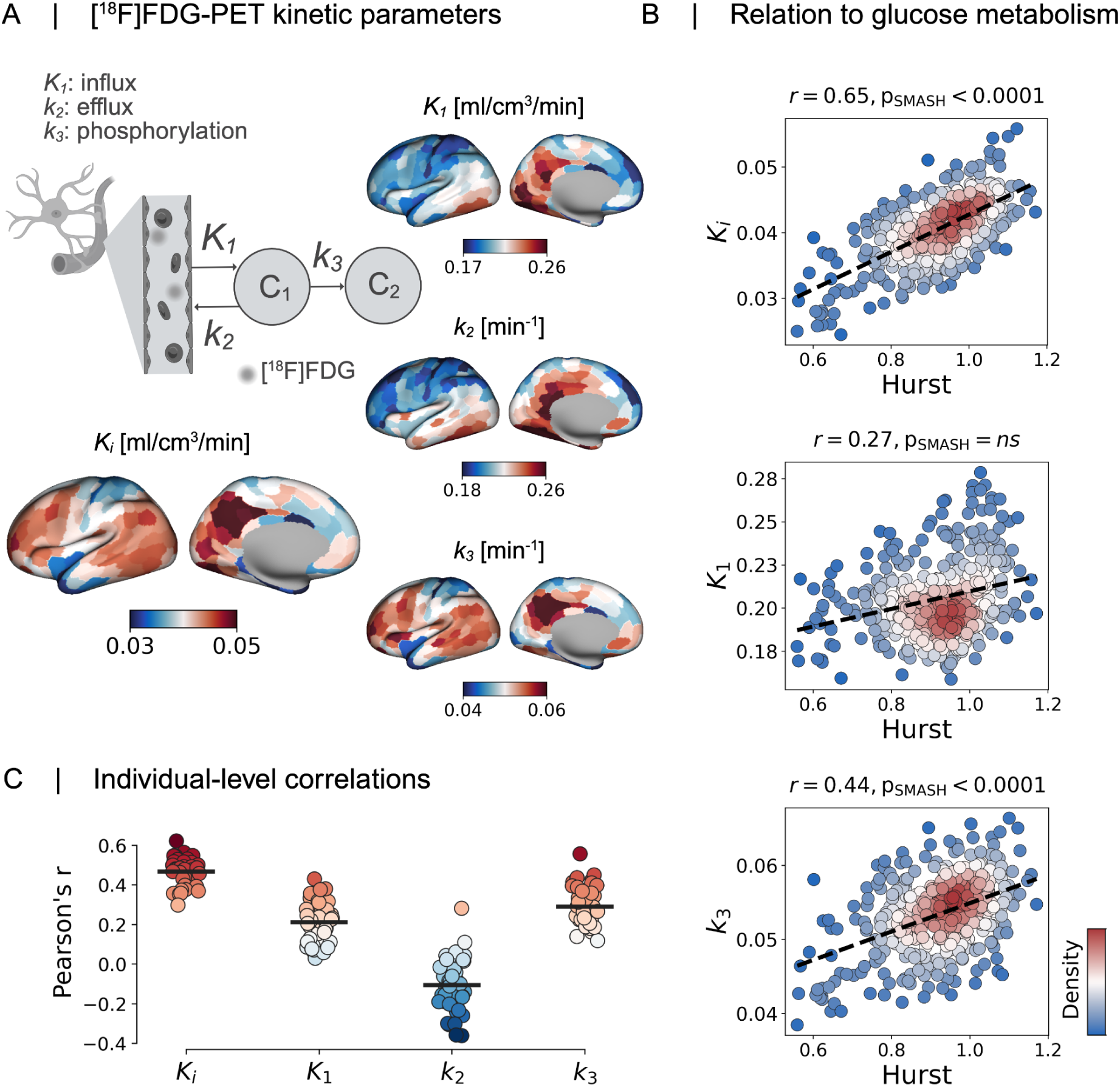
T**h**e **local cost of long-range temporal correlations.** A) Kinetic parameters describing [^18^F]FDG behavior: *K_1_*(tracer influx across the blood-brain barrier), *k_2_* (efflux or clearance of [^18^F]FDG into venous blood), and *k_3_* (phosphorylation by hexokinase). Additionally, the macro-parameter *irreversible uptake rate* (*K_i_*) is considered to describe the overall behavior. B) Group-level associations between the Hurst exponent and [^18^F]FDG kinetic parameters. Long-range temporal correlations were strongly associated with the macro-parameter *K_i_*. Looking at the micro-parameters, the strongest association was found with *k_3_*, i.e., phosphorylation by hexokinase. Correlation with *k_2_* is provided in the Supplementary Materials, Figure S3. C) Individual-level correlations between parameters and the Hurst exponent broadly recapitulate the group-level findings.

### Long-range temporal correlations and glucose metabolism are coupled across subjects

While spatial overlap between maps offers an intuitive, at-a-glance indication of their coupling, more compelling evidence emerges from across-subject associations. In other words, if at least a fraction of inter-individual differences in glucose consumption can be traced back to inter-individual differences in long-range temporal correlations, this would indicate a strong tethering of glucose metabolism to long-range temporal dependencies. Here, we hypothesize that brains characterized by stronger long-range temporal correlations (i.e., a higher average Hurst exponent) incur higher energetic costs than those with weaker correlations. To test this prediction, we linearly regress the Hurst exponent onto each [^18^F]FDG kinetic parameter, while including age, sex, and overall in-scanner motion as covariates. We identify a significant across-subject positive relationship between the Hurst exponent and [^18^F]FDG phosphorylation rate, indexed by *k_3_* (model *R^2^* = 0.20; *β* coefficient *t* = 3.28, *p* = 0.002; Figure 3). This finding suggests that brains exhibiting stronger long-range temporal correlations undergo a faster phosphorylation rate by hexokinase, i.e., higher glucose utilization. Importantly, the removal of the global signal does not impact the relationship between average Hurst exponent and *k_3_* (R^2^ = 0.23, *β* coefficient *t* = 3.46, *p* = 0.001). Collectively, these results underscore a relationship between persistent temporal dependencies and glucose consumption that goes beyond spatial similarity.

**Figure 3.**
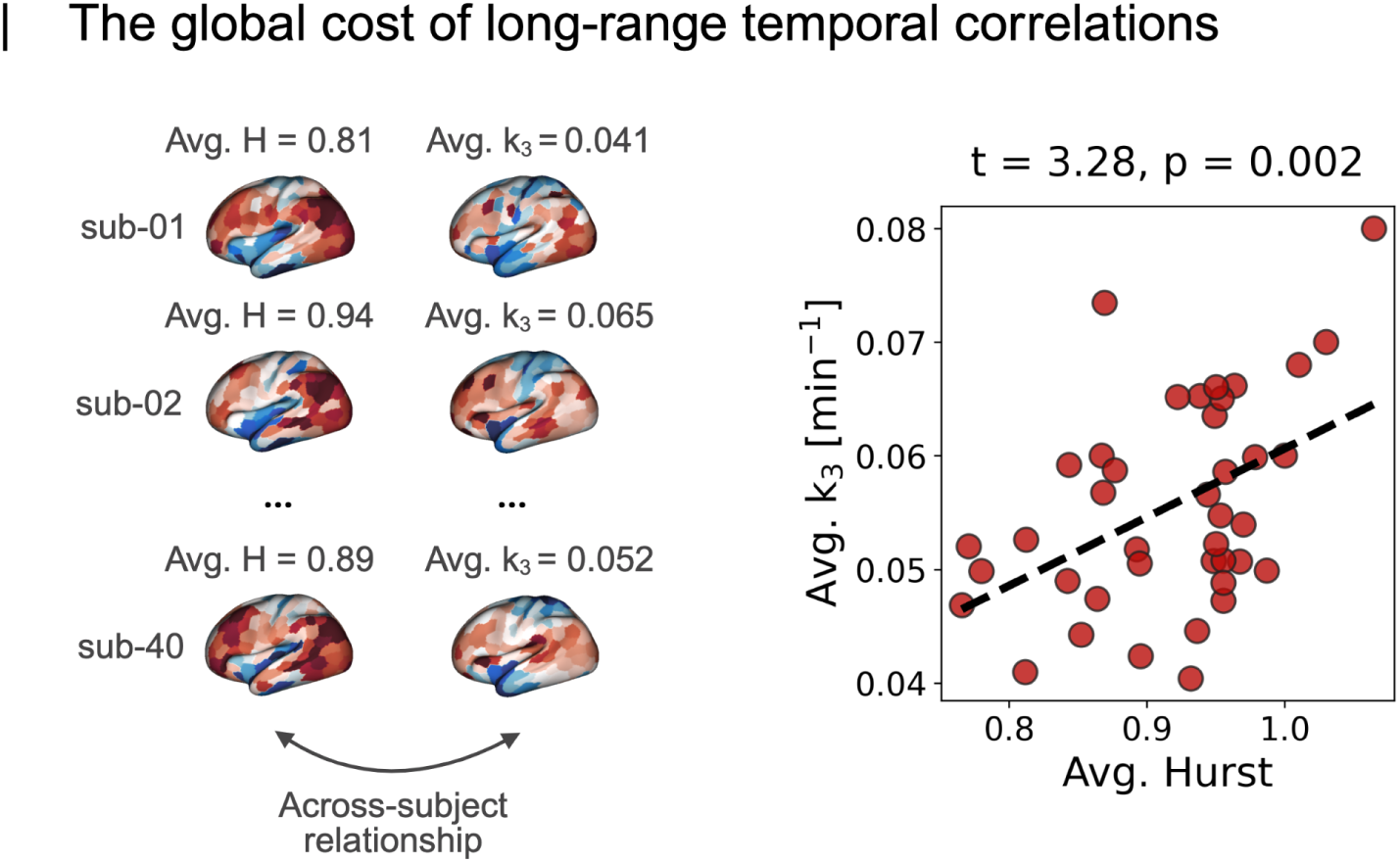
T**h**e **global cost of persistent temporal dependencies.** Across-subject relationship between average Hurst exponent and average k_3_. Each dot represents one subject from the experimental cohort. A higher average Hurst exponent is associated with a faster phosphorylation by hexokinase, i.e., higher glucose utilization. The regression model included age, sex, and in-scanner head motion as covariates, and explains a significant portion of the variance (*R^2^* = 0.20).

### Sustaining long-range temporal correlations requires both glucose and protein synthesis

We have shown the costly nature of long-range temporal correlations in terms of glucose metabolism. We now nuance the analysis by considering two additional biological properties: synaptic density and protein synthesis. This analysis is motivated by glucose consumption being both a process in itself and a source of fuel for other processes, including synaptic plasticity. Synaptic density here is based on [^11^C]UCB-J PET (Johansen et al., 2024), whereas the protein synthesis rate is derived from L-[1-^11^C]Leucine PET (Picchioni et al., 2023). The two template maps are based on independent data sources. We develop a simple multilinear regression model in which the Hurst exponent is regressed onto glucose metabolism (*K_i_*), synaptic density, and protein synthesis rate (Figure 4A). The model explains 67% of the variance, and its accuracy cannot be accounted for by confounding factors such as spatial autocorrelation (p_SMASH_ < 0.0001). Our results show that the Hurst exponent pattern is influenced by both energy use and protein production, with positive effects from glucose metabolism and protein synthesis, and a smaller negative effect from synaptic density. We then evaluate the importance of each predictor via dominance analysis, considering general dominance as the measure of interest (Budescu 1993). Protein synthesis rate shows the highest general dominance, making it the best predictor of long-range temporal correlations (Figure 4B). Glucose metabolism is the second-best predictor, with a general dominance of 25%. Net of other predictors, synaptic density does a poor job in explaining the regional patterning of the Hurst exponent. As a final step, we cross-validate our model using a distance-dependent approach (Hansen et al. 2021). For each node in the parcellation atlas, we train the multilinear regression model on 80% of its closest neighbors in Euclidean space and test it on the remaining 20% (Figure 4C). The model generalizes well to unseen, distant data, with an average correlation coefficient of r = 0.70 (SD = 0.02; Figure 4C). Taken together, these findings indicate that the long-term memory of neural activity may be supported by greater protein turnover. In other words, a faster rate of protein synthesis may be required to sustain a slower decay in the brain’s autocorrelation. Importantly, both glucose metabolism and protein synthesis exert independent effects on the Hurst exponent.

**Figure 4.**
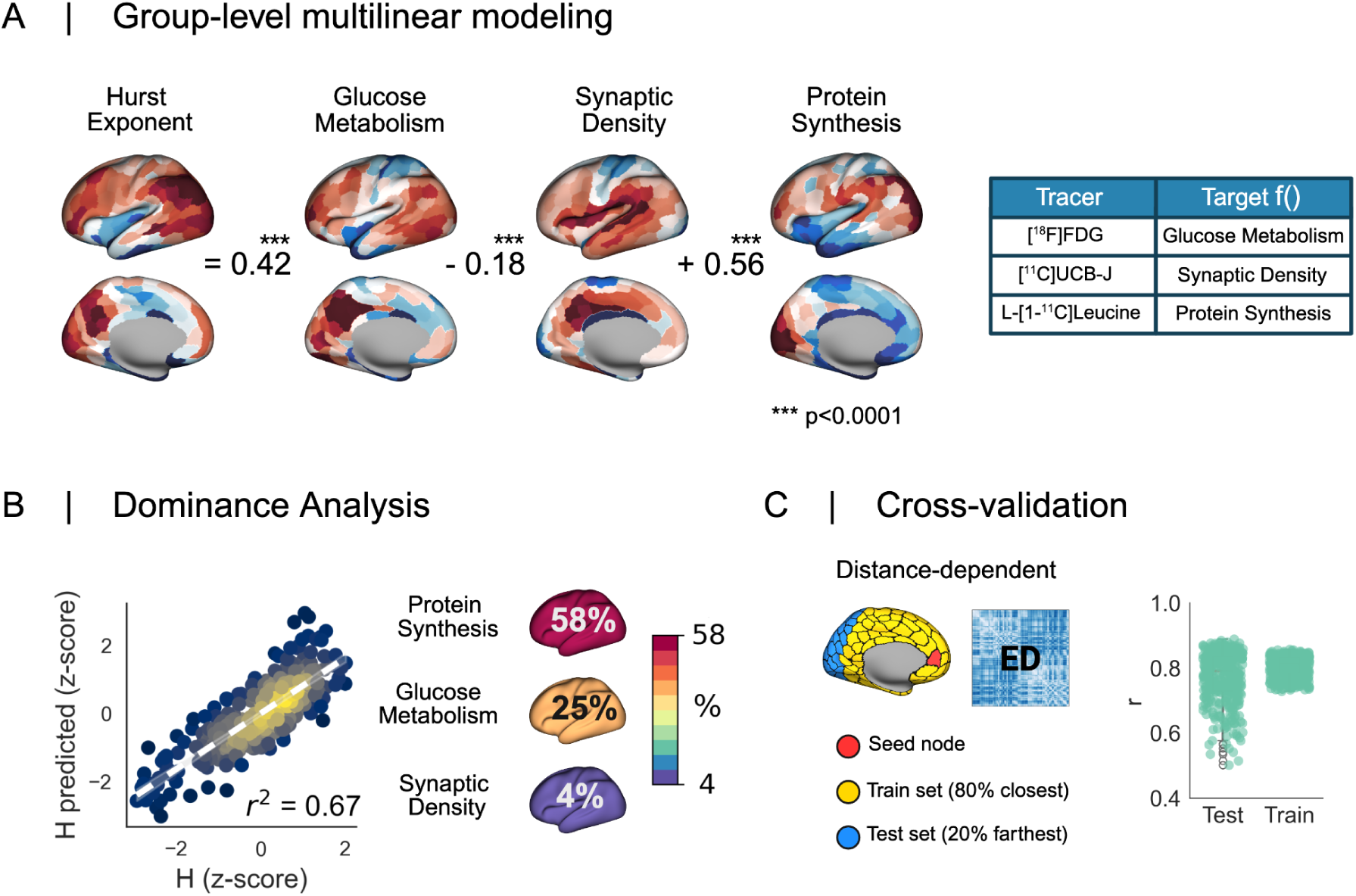
L**o**ng**-range temporal correlations require not only glucose but also consistent protein synthesis.** A) The group consensus Hurst exponent map is regressed onto three different aspects of brain biology: glucose metabolism, synaptic density, and rate of protein synthesis. The model *R^2^* is greater than that of null models that preserve spatial autocorrelation (p_SMASH_<0.0001). The reported beta coefficients are standardized. B) Dominance analysis for predictor importance. Protein synthesis rate had the highest dominance (58%), followed by glucose metabolism (25%). Synaptic density had little contribution in explaining the regional variation of the Hurst exponent. C) Distance-dependent cross-validation of the multilinear regression model (Hansen et al. 2021). For each node of the parcellation, the model was trained on 80% of the closest nodes (based on Euclidean distance) and tested on the farthest 20%. Model accuracy was assessed as the correlation coefficient. The average *r* in the test set was 0.70, confirming the validity of the model. Please note that, in this analysis, only cortical ROIs are considered, since the rate of protein synthesis is provided in surface space (see Methods section). ED indicates Euclidean distance.

## Discussion

In this multimodal effort, we sought to untangle the interplay between the brain’s long-range temporal correlations, indexed by the Hurst exponent, and glucose consumption. We exploited a unique dataset comprising rs-fMRI and [^18^F]FDG-PET data acquired in healthy subjects. The multimodal coupling between glucose metabolism and the Hurst exponent was assessed across multiple levels of analysis. We began with group-level associations and progressively incorporated individual-level data, thereby adding nuance to the systematic coupling. Lastly, we contextualized the Hurst exponent in light of two additional PET tracers, [^11^C]UCB-J and L-[1-^11^C]Leucine, used to map synaptic density (Finnema et al. 2016; Johansen et al. 2024) and the rate of protein synthesis (Schmidt et al., 2005; Smith et al. 2005), respectively.

Our results show that long-range temporal correlations are a pervasive feature of spontaneous brain activity, consistently observed across regions and individuals. This ubiquity suggests that scale-invariant neural dynamics are a hallmark of healthy brain function, as predicted by the Critical Brain Hypothesis (He 2014; Hesse & Gross 2014; Kitzbichler et al. 2009). According to this framework, operating near criticality enables the brain to strike an optimal balance between stability and flexibility, with long-range correlations reflecting the system’s capacity for long-memory processes and efficient information transmission (Cocchi et al. 2017; Expert et al. 2010). Our findings build on this framework by demonstrating a systematic relationship with glucose metabolism: the degree to which brain regions “remember” their past strongly predicts how much glucose they consume as fuel. Crucially, this relationship holds at both the group and individual levels. We then refined our analysis by examining across-subject relationships between the Hurst exponent and metabolic demand. Individuals exhibiting more pronounced long-range temporal correlations consistently displayed accelerated phosphorylation rates of [^18^F]FDG by hexokinase – i.e., increased metabolic utilization – as indexed by elevated *k_3_* values. Remarkably, interindividual variability in the Hurst exponent accounted for a significant fraction of the variance in [^18^F]FDG phosphorylation rates across participants. This underscores the central role of intrinsic neural dynamics in shaping the brain’s energetic profile. In essence, brains characterized by more temporally persistent activity patterns are associated with higher metabolic costs, suggesting that the self-similar, scale-free architecture of spontaneous brain activity – while enabling efficient information processing – exacts a significant energetic toll.

To better understand the origin of this metabolic burden, it is essential to consider the brain’s inherently networked organization (Bassett & Sporns 2017; Barabási et al. 2023; Bullmore & Sporns 2009; Sporns et al. 2005). Brain regions do not operate in isolation; rather, they function as parts of distributed, large-scale circuits. These interactions are energetically expensive, yet the associated cost is markedly asymmetric: up to 75% of the brain’s glucose consumption is attributed to postsynaptic activity (Attwell & Laughlin 2001; Harris et al. 2012; Riedl et al. 2016). This implies that metabolically demanding regions are likely to act as integrative hubs – nodes that receive and process large volumes of afferent input. Strikingly, evidence from the mouse brain shows that such receiver regions, characterized by dense afferent projections, tend to exhibit slower and more persistent temporal dynamics (Sethi et al. 2017). In concert, these findings suggest that the metabolic cost of maintaining long-range temporal correlations may reflect the integrative demands placed on afferent hubs, which must reconcile multisensory inputs – each with distinct temporal signatures – to support unified brain function (Gollo et al. 2015).

Beyond glucose consumption, our multimodal analysis reveals that the maintenance of long-range temporal correlations may also depend on sustained biosynthetic processes. This conclusion stems from the inclusion of an additional PET-derived marker of protein synthesis rate in our analysis (Bishu et al. 2008, Hellyer et al. 2017, Schmidt et al. 2005, Smith et al. 2005). While glucose fuels moment-to-moment neural activity, it also supports the biosynthetic machinery required to preserve neural structure and function over time. Our findings indicate that both glucose metabolism and protein synthesis contribute independently to the spatial distribution of long-range temporal correlations. These results suggest that sustaining persistent, long-memory dynamics is not only metabolically demanding but also biosynthetically intensive. In other words, the brain’s capacity to maintain temporally extended, scale-free fluctuations may rely on ongoing protein synthesis – potentially reflecting the continuous remodeling of synaptic and molecular scaffolds that support functional integration. This idea ties into a broader, overarching theme emerging from our findings: the brain’s ability to sustain slow, scale-invariant dynamics appears to hinge on fast, energy-and resource-intensive molecular processes.

Taken together, our results open new avenues for future research. For instance, if long-range temporal correlations are metabolically costly phenomena reflecting the brain’s proximity to criticality, a natural prediction is that conditions disrupting criticality – such as loss of consciousness – should exhibit regional reductions in long-range temporal correlations, accompanied by decreases in metabolic activity. Conversely, we hypothesize that impairments in glucose metabolism through aging, drugs, or disease will have specific effects upon long-range temporal correlations in brain activity. Intriguingly, recent theoretical work suggests that maintaining neural population activity in a “critical” state may help to balance energy costs and computational efficacy (Tatsukawa et al. 2025); future research will be needed to address how this balance is adjudicated at the meso-and macroscopic levels. Further, although our analyses focused on the resting state, future investigations should examine how the coupling between long-range temporal correlations and metabolism reconfigures during task-evoked states. Oxidative phosphorylation is the brain’s primary and most efficient pathway for ATP production (Rolfe & Brown 1997). However, to accommodate rapid surges in local energy demand, the brain can transiently increase its reliance on aerobic glycolysis – for reasons that remain unclear (Fox et al. 1988; Vaishnavi et al. 2010; Goyal et al. 2014). It is plausible that task-induced modulations of the Hurst exponent may be, at least in part, supported by localized increases in aerobic glycolysis, enabling dynamic adjustments to ongoing computational demands. Unraveling the balance between metabolism and the complex spatiotemporal organization of neural activity represents a compelling path toward understanding the energetic underpinnings of maintaining optimal local and large-scale brain dynamics.

## Methods

### Dataset

#### Demographics

The study cohort consists of 43 healthy subjects (22 female; mean age = 56.2 ± 13.4 years) drawn from the Adult Metabolism & Brain Resilience (AMBR) study (Goyal et al. 2023). All imaging procedures were approved by the Human Research Protection Office and the Radioactive Drug Research Committee at Washington University in St. Louis.

#### MR protocol

MRI data were acquired using a Siemens Magnetom Prisma^fit^ scanner. Each participant underwent a structural MRI scan using a 3D sagittal T1-weighted magnetization-prepared rapid gradient echo (MPRAGE) sequence with 180° radio-frequency pulses (multi-echo: TE = 1.81, 3.6, 5.39, 7.18 ms; TR = 2,500 ms; TI = 1,000 ms; voxel size = 0.8×0.8×0.8 mm). The average of the first two echoes was used to generate the final T1-weighted image. Following this, two sessions of T2*-weighted gradient-echo echo-planar imaging (GE-EPI) were collected (TR/TE = 800/33 ms, flip angle = 52°, voxel size = 2.4×2.4×2.4 mm, multiband factor = 6, 375 volumes), including additional spin-echo (SE) acquisitions (TR/TE = 6,000/60 ms, flip angle = 90°) for distortion correction.

#### PET protocol

Each participant underwent a single [^18^F]FDG-PET scan using a Siemens/CTI ECAT EXACT HR+ 962 PET scanner. Dynamic [^18^F]FDG-PET data were acquired for 60 minutes following the injection of 5.1 ± 0.3 mCi (187.7 ± 12.1 MBq) of [^18^F]FDG. Three 2-mL venous blood samples were collected during the scan to measure plasma radioactivity. Participants were instructed to keep their eyes closed while remaining awake throughout the PET acquisition. PET data were reconstructed using filtered back-projection with a ramp filter (5 mm FWHM). Attenuation correction was performed using a transmission scan. The dynamic reconstruction consisted of 52 frames of increasing duration: 24 × 5 s, 9 × 20 s, 10 × 1 min, and 9 × 5 min.

### Anatomical, rs-fMRI and PET processing

#### Structural MRI

The high-resolution T1-weighted image was bias-corrected, skull-stripped using SynthStrip (Hoopes et al. 2022), and processed with the standard surface reconstruction pipeline in FreeSurfer (version 7.4.1; Fischl 2013). All surface reconstructions and segmentations were visually inspected and manually corrected when necessary. Native surfaces were resampled onto the Conte69 surface using Connectome Workbench tools (Glasser et al. 2013). Registration to MNI space was performed using the deep-learning-based tool SynthMorph (Hoffmann et al. 2022), and the accuracy of each registration was visually checked.

#### rs-fMRI

Functional data were corrected for slice timing, head motion, and susceptibility-induced distortion using standard tools from the FMRIB Software Library (FSL, Jenkinson et al. 2012). Rigid registration of functional data to the high-resolution T1w was carried out using FreeSurfer’s *bbregister*. Minimally processed data were first projected onto each subject’s native surface and subsequently onto the Conte69 standard surface. Subcortical volumes were nonlinearly warped to 2 mm MNI space using the deformation fields previously estimated with SynthMorph. CIFTI files were generated by combining cortical surface vertices and subcortical voxels. Data were then high-pass filtered (cutoff: 0.008 Hz) and spatially smoothed with a 6 mm FWHM Gaussian kernel. Nuisance regression included the Volterra expansion of the six motion parameters (i.e., 24 regressors), as well as five principal components each from white matter and cerebrospinal fluid signals (aCompCor; Behzadi et al. 2007). Preprocessed data were parcellated using the Schaefer 400 regions 7 Networks atlas (Schaefer et al. 2018), augmented with the Tian subcortical atlas (Version S1, Tian et al. 2020).

#### PET

Dynamic [^18^F]FDG-PET data were first motion corrected using FSL’s *mcflirt.* Summing the PET frames from the interval between 40 and 60 min post-injection, a PET template map was obtained for registration purposes. Then, in order to obtain the input function for kinetic modeling, motion-corrected PET data underwent a semi-automatic pipeline to extract an image-derived input function (De Francisci et al. 2024). Three late venous samples were utilized to perform the Chen’s spillover correction (Chen et al. 1998). Data were then fitted to the two compartments Sokoloff model (Sokoloff et al. 1977) using a two-step procedure (Volpi et al. 2025). Firstly, k-means clustering was applied to voxel-level PET data to extract 6 Gray matter (GM) and 5 White matter (WM) clusters, on which the Sokoloff’s model was fitted using a weighted nonlinear least squares estimator. Secondly, cluster-wise estimates were propagated at the voxel level using a Variational Bayesian (VB) approach (Castellaro et al. 2017). Thus, for each subject the three parameters describing tracer kinetics were extracted: *K_1_* [mL/cm^3^/min] (influx), *k_2_* [min-1] (efflux), *k_3_* [min-1] (phosphorylation). Based on the three micro-parameters, the net *irreversible tracer uptake* (*K_i_*, [mL/cm^3^/min]) was simply obtained as a linear combination 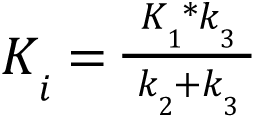 at the voxel level. All the estimated parameters were warped to standard space passing through the high-resolution T1w.

### Hurst exponent mapping

The Hurst exponent was estimated using a maximum likelihood approach within a fractionally integrated process (FIP) framework, implemented via Discrete Wavelet Transform (You et al. 2012). This method provides significant advantages over conventional fractional Gaussian noise (fGn) models by: (1) relaxing the stationarity assumption, and (2) relaxing the Hurst ≤ 1 constraint that frequently produces ceiling effects in neuroimaging data. The analysis was implemented using the nonfractal MATLAB toolbox (https://github.com/wonsang/nonfractal) and setting the following parameters: filter =’haar’, ub = [1.5, 10] and lb = [−0.5, 0] in line with previous works (Fotiadis et al. 2023, Trakoshis et al. 2020). The full details of the estimation approach are available in the validation paper (You et al. 2012).

### Independent datasets

#### Synaptic Density

The atlas is provided by Johansen et al. (2024) and is publicly available online (https://xtra.nru.dk/SV2A-atlas/). Briefly, thirty-three healthy subjects underwent [^11^C]UCB-J PET, with continuous automatic arterial sampling (first 15 minutes) followed by manual sampling to obtain a metabolite-corrected arterial input function (AIF). Data were resampled onto standard surface (*fsaverage*) and volumetric (*MNI152*) spaces, followed by spatial smoothing (10 mm FWHM for surface data and 5 mm for volumetric data). Data were fitted with a one-tissue compartment model (1TCM) to obtain tissue distribution volumes (V_T_, [mL/cm³]) for each voxel/vertex. Ex-vivo [³H]UCB-J autoradiography was later used to convert V_T_ values to cerebral SV2A protein densities (B_max_, [pmol/mL]). For full methodological details we refer to the validation paper (Johansen et al. 2024). The atlas in fsaverage space was parcellated according to the Schaefer 400 regions 7 Networks atlas (Schaefer et al. 2018).

#### Cerebral Protein Synthesis Rate (rCPS)

The atlas is provided by Picchioni et al. (2023) and is publicly available in OpenNeuro (doi:10.18112/openneuro.ds004733.v1.0.0). In brief, 17 subjects underwent dynamic L-[1-^11^C]Leucine PET and arterial sampling to measure the concentration of labeled and unlabeled leucine in blood plasma. The parameters of the model describing L-[1-^11^C]Leucine kinetics were fitted using a basis function method (BFM) and a voxel-level map of rates protein synthesis (rCPS, [nmol/g/min]) was obtained. See the original papers for methodological details (Picchioni et al. 2023; Schmidt et al. 2005; Smith et al. 2005; Tomasi et al. 2009; Veronese et al. 2018). The atlas in fsaverage space was parcellated according to the Schaefer 400 regions 7 Networks atlas (Schaefer et al. 2018).

### Statistics and null models

#### Intraclass correlation coefficient (ICC)

We assessed reliability using the Intraclass Correlation Coefficient (ICC, McGraw & Wong, 1996). We used the ICC-(1,1) model for one-way random effects and single measurements per session. The ICC-(1,1) was implemented in Python using the Pingouin library (https://github.com/raphaelvallat/pingouin).

#### Spatial autocorrelation-preserving null models

Spatial autocorrelation can bias measures of statistical similarity between brain maps (Markello & Misic 2021). In this work, we mitigated this issue by constructing appropriate null models when assessing spatial associations. These null models (N = 10,000) were based on the variogram-matching approach developed by Burt et al. (2020) as implemented in *BrainSMASH* (https://github.com/murraylab/brainsmash). This approach is able to accommodate both cortical and subcortical structures. Critically, the agreement between empirical vs. surrogate variograms was ensured to validate the null model (supplementary materials, figure S2).

#### Linear regression for across-subject relationships

To investigate across-subject relationships between long-range temporal correlations and glucose metabolism, we averaged both the Hurst exponent and glucose parameters (*K_i_, K_1_, k_3_*) across brain regions. We then regressed the average Hurst exponent onto each of the [^18^F]FDG kinetic parameters separately. Only metabolically relevant parameters were considered; therefore, *k_2_* was excluded from the analysis. Each linear model included age, sex, and in-scanner head motion as covariates. Head motion was quantified using the average framewise displacement (FD). We assessed the significance of each predictor by examining the t-statistic associated with its regression coefficient. All models were fitted using the Python library *Scikit-learn* (https://github.com/scikit-learn/scikit-learn).

#### Linear regression model based on independent PET maps

We linearly regressed the group-averaged Hurst exponent map onto three measures derived from kinetic modeling of [^18^F]FDG (Ki, [mL/cm^3^/min]), [^11^C]UCB-J (Synaptic density, B_max_, [pmol/mL]), and L-[1-^11^C]Leucine (Cerebral Protein Synthesis rate, rCPS, [nmol/g/min]) PET data. Both predictors and the dependent variable were z-scored across brain regions prior to model fitting. This standardization removes the intercept and yields standardized beta coefficients. The model was fitted using Ordinary Least Squares (OLS), as implemented in *Scikit-learn* (https://github.com/scikit-learn/scikit-learn).

#### Dominance analysis

We conducted a dominance analysis to rank predictors according to their relative importance in explaining the regional distribution of the Hurst exponent. To this end, we regressed the Hurst exponent across brain areas onto each possible subset of predictors (2ᵖ–1 models, where *p* is the total number of predictors, here 3). We used *general dominance* as the measure of importance, defined as the average incremental contribution of each predictor to R² across all subset models (Budescu 1993). Dominance analysis was implemented using *netneurotools* (https://github.com/netneurolab/netneurotools).

#### Distance-dependent cross-validation

The multilinear model predicting the Hurst exponent from PET maps was cross-validated using a distance-dependent approach, originally developed by Hansen et al. (2021). Inter-nodal distances were computed as the Euclidean distance between the centroid coordinates of brain regions defined in the parcellation atlas. For each seed node, model coefficients were estimated using data from 80% of the regions closest to the seed. The model was then tested on the remaining 20% of the farthest regions. Prediction accuracy was assessed using Pearson’s correlation between the empirical and predicted Hurst exponent values in both training and test sets.

## Data and code availability statement

The data are available upon reasonable request. They are not publicly shared to protect the privacy of research participants. The code for estimating the Hurst exponent is publicly available in the MATLAB *nonfractal* toolbox (https://github.com/wonsang/nonfractal). Functions for ICC(1,1) and other versions are available in the Python package Pingouin (https://github.com/raphaelvallat/pingouin). The code for SA-preserving null models is available in the BrainSMASH package (https://github.com/murraylab/brainsmash). The function for dominance analysis is included in the Netneurotools toolbox (https://github.com/netneurolab/netneurotools).

## Author Contribution

**M.F.** Conceptualization, Methodology, Software, Formal analysis, Writing – Original Draft.

**A.R**. Software, Writing – Review & Editing. **M.C.** Writing – Review & Editing. **C.T.** Writing – Review & Editing. **I.M.** Writing – Review & Editing. **T.V.** Formal analysis, Writing – Review & Editing. **A.G.V.** Data curation, Writing – Review & Editing. **M.S.G**. Data curation, Writing – Review & Editing. **M.C.** Conceptualization, Writing – Review & Editing. **A.B.** Conceptualization, Methodology, Supervision, Writing – Review & Editing.

## Acknowledgements & Funding information

We are grateful to all research participants for their altruism in contributing to these data. We also thank the Neuroimaging Laboratory, the Knight Alzheimer’s Disease Research Center, and the cyclotron and imaging staff for their essential role in enabling the AMBR dataset collection.

This work was supported by the National Institutes of Health (NIH) under grants R01AG053503, R01AG057536, RF1AG073210, and RF1AG074992. Support for the research, authorship and/or publication of this article was provided by #NEXTGENERATIONEU (NGEU) and funded by the Ministry of University and Research (MUR), National Recovery and Resilience Plan (NRRP), Project MNESYS (PE0000006) - A Multiscale integrated approach to the study of the nervous system in health and disease (DN. 1553 11.10.2022).

