## Supplementary Materials for "Glucose Metabolism echoes Long-Range Temporal Correlations in the Human Brain"

| Hurst exponent subcortex

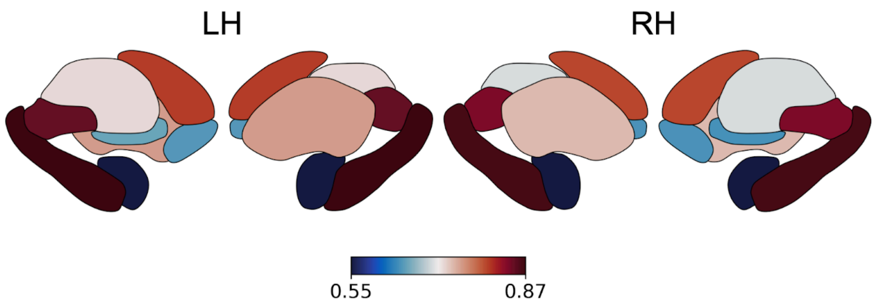


**Figure S1. Hurst exponent distribution across subcortical structures.** Subcortical data were parcellated using the Melbourne/Tian subcortical Atlas at its coarser granularity level (S1, 16 regions). As in the cortex, Hurst exponent values across subcortical structures were consistently above the 0.5 threshold, indicating persistent neuronal dynamics. The highest values were found in the hippocampus, and the lowest in the amygdala. For a full list of subcortical regions, see the validation paper by Tian et al. (2020).

| Null model (BrainSMASH) fit quality and examples

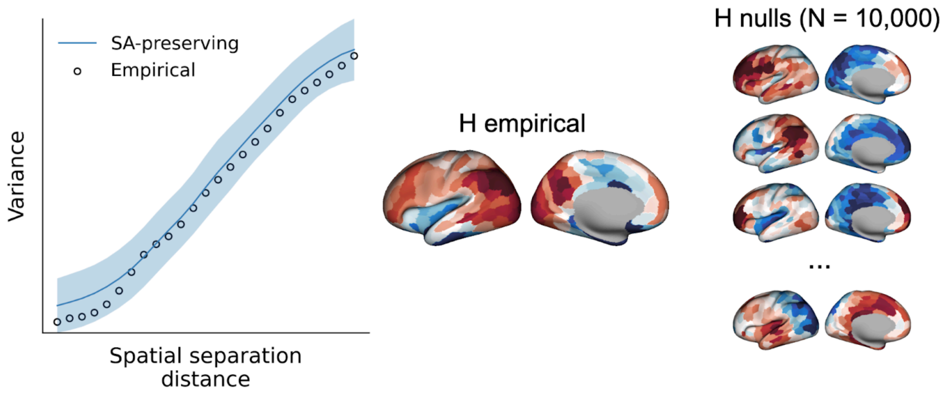


**Figure S2. Quality of the brainSMASH fit and examples of null maps.** All correlation coefficients and linear models in this study were tested against null distributions generated using BrainSMASH (Burt et al. 2020). The left panel shows the fit quality between empirical and surrogate variograms, confirming that the null maps preserve the spatial autocorrelation structure. The right panel shows representative null maps derived from the Hurst exponent map, each preserving the spatial autocorrelation of the empirical data.

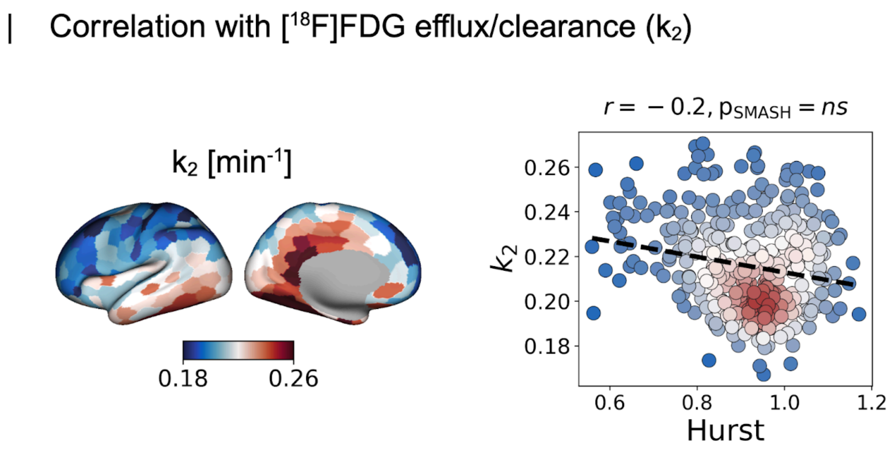


**Figure S3. Correlation between Hurst exponent and $k_2$ at the group level.** No significant correlation was observed between the Hurst exponent and the [$^{18}$F]FDG-PET microparameter $k_2$ [$min^{-1}$]. This parameter reflects the efflux or clearance of the tracer into venous blood and is therefore not directly informative about cellular metabolism.
